# Genome-wide haplotype association study in imaging genetics using whole-brain sulcal openings of 16,304 UK Biobank subjects

**DOI:** 10.1101/2020.08.26.267617

**Authors:** Slim Karkar, Claire Dandine-Roulland, Jean-François Mangin, Yann Le Guen, Cathy Philippe, Jean-François Deleuze, Morgane Pierre-Jean, Edith Le Floch, Vincent Frouin

**Affiliations:** Université Paris-Saclay, CEA, Neurospin, 91191, Gif-sur-Yvette, France; Université Paris-Saclay, CEA, Centre National de Recherche en Génomique Humaine, 91057, Evry, France

**Keywords:** Imaging-genetics study, Haplotype Association Analysis, UK Biobank

## Abstract

Neuroimaging-genetics cohorts gather two types of data: brain imaging and genetic data. They allow the discovery of associations between genetic variants and brain imaging features. They are invaluable resources to study the influence of genetics and environment in the brain features variance observed in normal and pathological populations. This study presents a genome wide haplotype analysis for 123 brain sulcus opening value (a measure of sulcal width) across the whole brain that include 16,304 subjects from UK Biobank. Using genetic maps, we defined 119,548 blocks of low recombination rate distributed along the 22 autosomal chromosomes, and analyzed 1,051,316 haplotypes. To test associations between haplotypes and complex traits, we designed three statistical approaches. Two of them use a model that includes all the haplotypes for a sin gle block, while the last approach considers one model by haplotype. All the statistics produced were assessed as rigorously as possible. Thanks to the rich imaging dataset at hand, we used resampling techniques to assess False Positive Rate for each statistical approach in a genome-wide and brain-wide context. The results on real data show that genome-wide haplotype analyses are more sensitive than single-SNP approach and account for local complex Linkage Disequilibrium (LD) structure, which makes genome-wide haplotype analysis an interesting and statistically sound alternative to the single-SNP counterpart.

## 1 Introduction

Numerous population-imaging studies have been built since the beginning of years 2000 based on earlier pioneering studies [1] to support researches mainly in vascular, neurodegenerative diseases or psychiatric syndromes [2] and now include genetics [3, 4, 5]. Our work is based on the UK Biobank resources [6, 7, 8], currently the most emblematic imaging-genetics cohort available as open-data. The UK Biobank cohort brings the unique opportunity to study the genetic and environmental dissection of numerous diseases or complex traits related to brain *via* imaging endophenotypes. These endophenotypes are structural or functional imaging-derived characteristics (or imaging-derived phenotypes - IDP [7] in UK Biobank) and DNA genotyping arrays provide the genetic measures [6]. In aging of normal and pathological brains, we observe the phenomenon of sulcal widening for which several related IDP like sulcal depth, sulcal opening and grey matter thickness in brain were shown by our group to be highly heritable [9]. In the case of grey matter thickness and sulcal opening, our group in [10] has also shown associations with several new markers using Genome-Wide Association Study (GWAS).

In order to yield robust inferences and interpretations from GWAS approaches, one should only consider the hits passing a strict genomic significance threshold [11] to account for the large number of tests. Moreover, in imaging-genetics studies, an additional correction for multiple phenotype testing is mandatory. After these corrections and in the case of a non-synonymous SNP hit, a straightforward interpretation can be carried out to hypothesise an association with the disease. In most cases, GWAS results are harder to interpret because the association is carried by a group of SNPs located in a non-coding region. In some cases, the grouping can be explained by a leading causal SNP which signal is spread amongst the neighbouring SNPs *via* Linkage Desequilibrium (LD). Using imputed SNPs, one can identify putative unmeasured causal variants. Nowadays more and more GWAS are using imputed SNPs to do fine mapping or allow replication study. An alternative to imputation is haplotype analysis that can also capture unmeasured variants.

One might also suspect that the association is actually carried by more than one SNP in the same region. In the case of multiple causal SNPs, burden-test or collapsing test have shown a high power to detect small effect sizes [12]. A gain in sensitivity can also be obtained by considering the combinations of alleles from several genotyped SNPs in the form of haplotypes [13].

Haplotypes in one individual consist in the combinations of several SNP alleles to form nucleotide sequences. These sequences of alleles can be obtained from phased SNPs that define which alleles are on the same chromosome and inherited together from each parent.

Both phased SNPs and imputed variants are available in the UK Biobank resources, based on large reference panels (1000 Genomes and the Haplotype Reference Consortium (HRC), see [6]). The high quality of the phasing dataset (assessed using trios of parents-child) is an incentive to use this information in haplotype-based analyses. The method described in this paper differs from the commonly used haplotype-based approaches on several points. First it applies at the genome level. Second, the haplotype-based tests are based on phased data which is unusual: out of the 9 haplotype-based methods cited in [14], only one (WHaIT, [15]) uses phase information as input. Moreover our method identifies individual haplotype associations by testing all haplotypes in a given genomic interval, while the WHaIT method only performs a global test, i.e. it does not identify which individual haplotype in the block is associated with the phenotype. Also of note is that the WHaIT method is only designed for categorical traits.

In this work, we propose to push the genome-wide haplotype association approaches to fit the specificity of the IDP obtained in imaging-genetics, more precisely sulcal opening measurements derived from the UK Biobank high quality imaging data. The aim of the paper is three-fold. First, we present this set of quantitative IDP and a normalization of their distributions across the subjects. We show their spatial variability which reveals that IDP are many and varied, and which motivates our search for even more sensitive association methods to compensate for the multiple testing issue. Second, we detail three tests for genome-wide haplotype associations with the traits. This part includes definition of blocks along the genome based on a genetic map, which is a prerequisite to any haplotype-based test definition. Third, we compare association hits obtained from these three haplotype-based tests and also with the regular genome-wide association test based on single SNPs. Finally, we carried out phenotype permutations to check the validity of our tests on the whole genome.

## 2 Material and Method

### 2.1 Samples

The present analyses were conducted under UK Biobank data application number 25251. The UK Biobank is a health research resource including genotypic data of about 500,000 people aged between 45 and 73 years old, that were recruited in the general population across United Kingdom.

The UK Biobank is expected to provide multi-modal MR brain images in 100,000 participants and, by March 2018 it had 20,060 subjects with a T1-weighted MRI [8]. These data were processed locally through BrainVisa/Morphologist pipeline [16] and quality controlled yielding a set of labelled cortical sulci for each of 18,175 subjects (see [10] for details).

We relied on the Quality Control carried out by the UK Biobank consortium on the genotyping data which excluded individuals with high missingness, high heterozygosity, first degree related individuals or sex mismatches [6]. With 658, 720 phased SNPs (Haplotype dataset), 784, 256 genotyped SNPs and 93, 095, 623 imputed variants, 16, 304 subjects passed the image processing and the genetic QC (48% of males and 52% of females).

### 2.2 Data Processing

#### Imaging data, sulcal opening

For each subject, 123 labelled sulci were extracted from T1-weighted images. For each sulcus, a measure of its width - a feature called opening - is computed as the ratio of the volume of the Cerebrospinal Fluid the sulcus contains to the surface of the sulcus (Figure 1).

**Figure 1:**
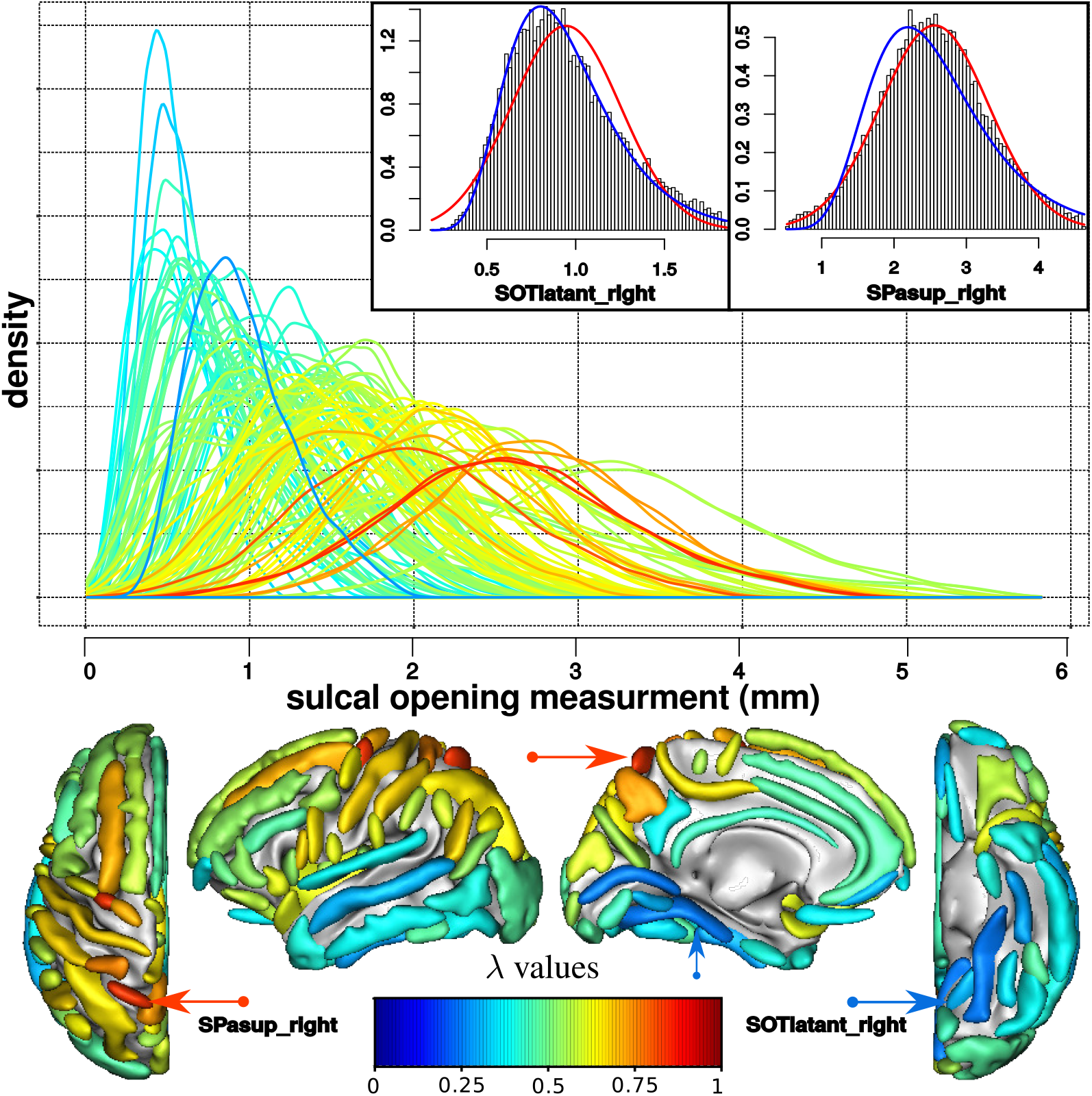
Distributions of sulcal opening measurements of 16, 304 subjects. In a sample of 16,304 subjects, sulcal opening measurements have different distributions across the brain, from normal distribution (red, λ = 1) to log-normal (blue, λ = 0). (Top) Distributions of uncorrected sulcal opening measurements, for 123 sulci, coloured according to λ values. Inlet panels are histograms of opening values for two sulci with fitted curves superimposed: (left) Anterior occipito-temporal sulcal opening distribution is almost log-normal (blue line); (right) Superior parietal sulcal opening distribution is almost normal (red line). (Bottom) Spatial map of λ values for sulci of the left hemisphere, coloured according to the λ value of their distribution. From left to right: top view, interior view, parietal view and bottom view.

For each sulcus, after adjusting for age and sex using linear regression, we identified and excluded outliers in the residual distribution using the robust interquartile range (IQR) method [17].

Then, since the distributions of sulcal opening values could exhibit deviation from the normal distribution, we evaluated this deviation and normalized them using a one-parameter Box-Cox transformation (power transformation) [18], see Section Supplementary Material 1. We selected the optimal power λ using goodness-of-fit for normal distribution (available in MASS R package) for each sulcus. The value of λ quantifies the similarity with a normal distribution. For a sulcus with an original normal distribution, λ =1 and for a sulcus with an original log-normal distribution, λ = 0.

In the following, the Box-Cox transformations of the sulcal openings, 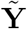, were considered as phenotypes.

#### Definition of Haplotypes blocks in UK Biobank

When a group of SNPs are in linkage disequilibrium (LD), they are not inherited independently: the presence/absence of one allele at one locus depends on the alleles of neighbouring loci in LD. Using allele frequencies and LD information available in reference panels, phasing softwares like SHAPEIT [19] estimate for each chromosome the sequence of SNP alleles that have been inherited together from each parent.

Using this approach, the UK Biobank consortium has released a curated genomewide dataset of 658, 720 phased SNPs over the 22 autosomal chromosomes [6]. From these phased data we define haplotypes, which are combinations of neighbouring SNP alleles on a single chromosome. We propose the following two-fold procedure to ensure the correct haplotype estimation.

First, we determine haplotype blocks that includes adjacent SNPs in high LD or in other words, present a low recombination rate. We used the GRCH37 Rutgers genetic map [20] that includes positions in base pairs (bp) and centiMorgans (cM) based on the 1000 Genomes project. For variants of the UK Biobank arrays that are not present in the genetic map, the position in cM was estimated using linear interpolation based on the local recombination rate (in cM/bp) within the interval defined by the two closest neighbouring variants in the map. For first (resp. last) assessed variants on a chromosome that fall outside the genetic map, recombination rates were estimated by linear extrapolation on the whole chromosome. The complete genetic map allows to define non-overlapping haplotype blocks in which any two consecutive SNP loci are less distant than *δ* cM. A distance greater than δ cM defines the start of a new block. Second, using the phased SNPs dataset and the blocks described above, we determine the haplotypes.

To insure an homogeneous cover of the genome, we chose δ value equals to 0.001 cM, leading to 119,548 haplotype blocks across the whole genome (see Supplementary Material 4 and [21])

### 2.3 Genome-wide haplotype association

#### 2.3.1 Count matrix of each haplotype block

For each haplotype block obtained previously, we defined the reference haplotype *h*_0_ as the most common one, and the *count matrix*, **H** = [**h_i_** … **h_m_**] ∈ {0, 1, 2}^(*N*×*m*)^, with *N* the sample size and *m* the number of alternative haplotypes denoted by *h*_1_,…, *h_m_*. Each element *h_i,j_* of **H** corresponds to the number of copies of the alternative haplotype *h_j_*, 1 ≤ *j* ≤ *m* for the individual *i*, 1 ≤ *i* ≤ *N*. In this way, the count matrix **H** codes for an additive haplotype model - the effect of each haplotype block corresponds to the sum of the effects of observed alternative haplotypes. Section Supplementary Material 2 shows an example of matrix **H**, obtained with 3 phased SNPs and 3 subjects.

#### 2.3.2 Association tests

We defined three association tests between haplotypes *h_j_* included in matrix **H**, and the phenotypes 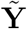, the sulcal opening values obtained after Box-Cox transformation as described previously. In the following, **X** denotes the matrix of covariates containing age, sex and the first ten principal components provided by UK Biobank to account for population stratification. Although all presented methods are using similar statistics, the three tests are addressing three different questions.

##### Haplotype block model test

In this test, referred as “block-test”, the following linear model is considered:

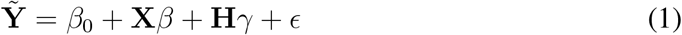

with 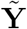 the phenotype vector, **X** the matrix of covariates and **H** the matrix of the m alternative haplotypes, included with fixed effects *β* and *γ* respectively, and *ϵ* the error vector. *β*_0_ is the intercept containing the effect of the reference haplotype.

This first test assesses the association between a given block, defined as the set of its haplotypes, and each phenotype i.e. *H*_0_: *γ* = **0_m_** versus *H*_1_: *γ*_1_ ≠ 0 or … or *γ_m_* ≠ 0

The significance of the association is estimated using a total variance test for nested linear model.

##### Complete model haplotype test

The second test is a complete model haplotype test and is referred as “complete-test”. It aims to test the association between one phenotype and each haplotype *h_j_* inside the block, while considering in the model the other haplotypes of the block. For this purpose, we used the same linear model regression as in the block-test (eq. 1) and we considered for each haplotype *h_j_*, 1 ≤ *j* ≤ *m*, the null hypothesis: *H*_0_: *γ_j_* = 0 versus *H*_1_: *γ_j_* = 0. For this test, we computed a two-sided *p*-value of the *t*-statistic.

##### Single haplotype model test

To test the effect of each haplotype versus the others, the third test referred as “single-test” is based on the following linear model:

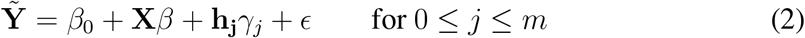

with 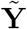 the phenotype vector, **X** the matrix of covariates and **h_j_** the count vector for the haplotype *h_j_*, included with fixed effects *β* and *γ*_j_ respectively, and *ϵ* the error vector. *β*_0_ is the intercept containing the mean effect of all haplotypes excluding the considered haplotype *j*. As previously, we computed a standard two-sided *p*-value of the *t*-statistic for the null hypothesis *H*_0_: *γ_j_* = 0 versus *H*_1_: *Y_j_* ≠ 0.

All three tests were implemented in R v.3.4.0 to run on a distributed HPC (code available upon request).

#### 2.3.3 Comparison with single-SNP association used in classical GWAS

To our knowledge, a comprehensive power analysis of haplotype association for quantitative traits on large cohorts (more than 10,000 subjects) has not been done. To gain insight into the power of the haplotype association tests, we propose a comparative study, where we match the results of haplotype-based tests with the results from the single SNP association used in classical GWAS considering both genotyped and imputed SNPs. The test used in classical GWAS using linear model in PLINK [22] is described in Section Supplementary Material 3

#### 2.3.4 False Positive Rates in the Genome-wide haplotype association tests

In order to evaluate the validity of the different haplotype association tests scrutinized in this study, we constructed datasets under the null hypothesis using permuted phenotypes. Then, we computed the *p*-values distributions and the associated False Positive Rate (FPR). The FPR is the percentage of tests that show a significant association in the permuted dataset at the significance threshold with the Bonferroni correction. We fixed the significance threshold to *α_N_t__* = *α*/*N_t_*, with *α* = 0.05, and *N_t_* is the number of hypotheses tested.

Correlations between variables are present in genetic and imaging datasets; between parts of the genome and within the brain respectively. To quantify the impact of these correlations, we computed the FPR under null hypothesis for several scenarios that keep or not the correlation structures among haplotype blocks and among phenotypes.

##### False Positive Rate while preserving the correlation within each haplotype block

In the first permutation scenario, a phenotype is randomly permuted for each block. Therefore, correlations between haplotype blocks across the genome present in the original dataset were removed in this scenario, preserving only the correlation within each haplotype block. The complete-test and the single-test produced several statistics per haplotype block (depending on the number of haplotypes), while the block-test produced a unique statistic per block. We computed *FPR* = *N_FP_/N_t,p_*, where *N_FP_* is the observed number of false positives and *N_t,p_* is the number of statistics produced by each of the three tests for the permuted phenotype *p*.

This permutation scenario was replicated on three phenotypes of sulcal opening: two that represent the range of λ values and one for the highest association found in previous study [10], with λ values ≈ 0.2, ≈ 0.8 and ≈ 0.6 respectively. The same permutation order was applied to the three phenotypes.

##### False Positive Rate while preserving the correlation within the genome

In the second permutation scenario, we used all the phenotypes and produced a permutated dataset where the phenotypes were randomly shuffled with the same permutation for all haplotype blocks along the genome.For each of the three tests, we considered all the statistics produced and computed *FPR_p_* = *N_FP_/N_t,p_* where *N_t,p_* is the number of statistics produced for a given permutated phenotype *p*. This analysis produced three FPRs per permuted phenotype that preserves the structure of the correlation within the genome. Reporting FPRs for each phenotype will enable the detection of phenotypes with low quality measurements.

This procedure was replicated ten times. To reduce the computational burden in this case, we considered the residuals 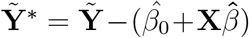, that is, the phenotypes 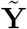 adjusted for the covariates **X**.

##### False Positive Rate while preserving the correlation within the genome and among the phenotypes

Last, we used the above described permutated dataset and, instead of considering each phenotype independently, we pooled the results across the phenotypes. For each permutation and each haplotype test, we considered all the statistics produced. We computed *FPR_T_* = *N_FP_/N_T_* where *N_T_* is the number of statistics produced by each test *for all permutated phenotypes*. This analysis produced three FPRs per permutation, that preserve both the structure of the correlation within the genome as well as the correlation among phenotypes.

## 3 Results

In this part, we present results obtained on the original data: the opening measurements of 123 brain sulci and genome-wide haplotypes of 16, 304 subjects. First, we present the effects of normalization of the phenotype distributions, and the characteristics of the haplotypes generated with the original data. Second, we present the significant association hits obtained with our genome-wide haplotype association study and provide the genomic location and length of significant haplotype blocks. Finally, we compare the differences in sensitivity on original data for the three tests proposed above and we propose a comparison study with the single-SNP association test.

A study of the False Positive Rate is also presented, where we study the FPR for the three models under the null hypothesis, using permutated datasets as described in Section 2.3.4.

### Imaging data processing

The distributions of sulcal opening measurements across the brain exhibit a large range of density shapes from quasi-normal distributions (max. λ = 0.8) to log-normal distributions (min λ = 0.2). Fig. 1 displays the λ parameter associated with the opening distribution for each sulcus across the brain.

We observed a pattern, with larger (in terms of median value) and more normally distributed sulcal opening measurements in parietal and occipital lobes. Conversely, we observed narrower, more log-normally distributed opening values for lower and inferior temporal sulci. This could be related to the differential aging rate of brain structures, such as the widening sulci rates that might differ across the brain [23].

### Genomic data processing

Across the 22 autosomal chromosomes, we tested over 1 million haplotypes from 119,548 blocks for association with the opening of 123 sulci. Candidate haplotypes had a length, on the genetic map, ranging from 1.4 × 10^-8^ to 1.7 × 10^-2^ cM (see Fig. 2, right panel), with a median value of 6 × 10^-4^ cM, corresponding to consecutive SNP runs of length ranging from 2 to 64. Left panel of Fig. 2 shows the coverage of the blocks along the 22 autosomal chromosomes. Details regarding the distribution can be found in the Section Supplementary Material 4. Of note is that the variation of the block length across the chromosomes remains small as it can be seen from the curve on the right of Fig 2 (similar distributions).

**Figure 2:**
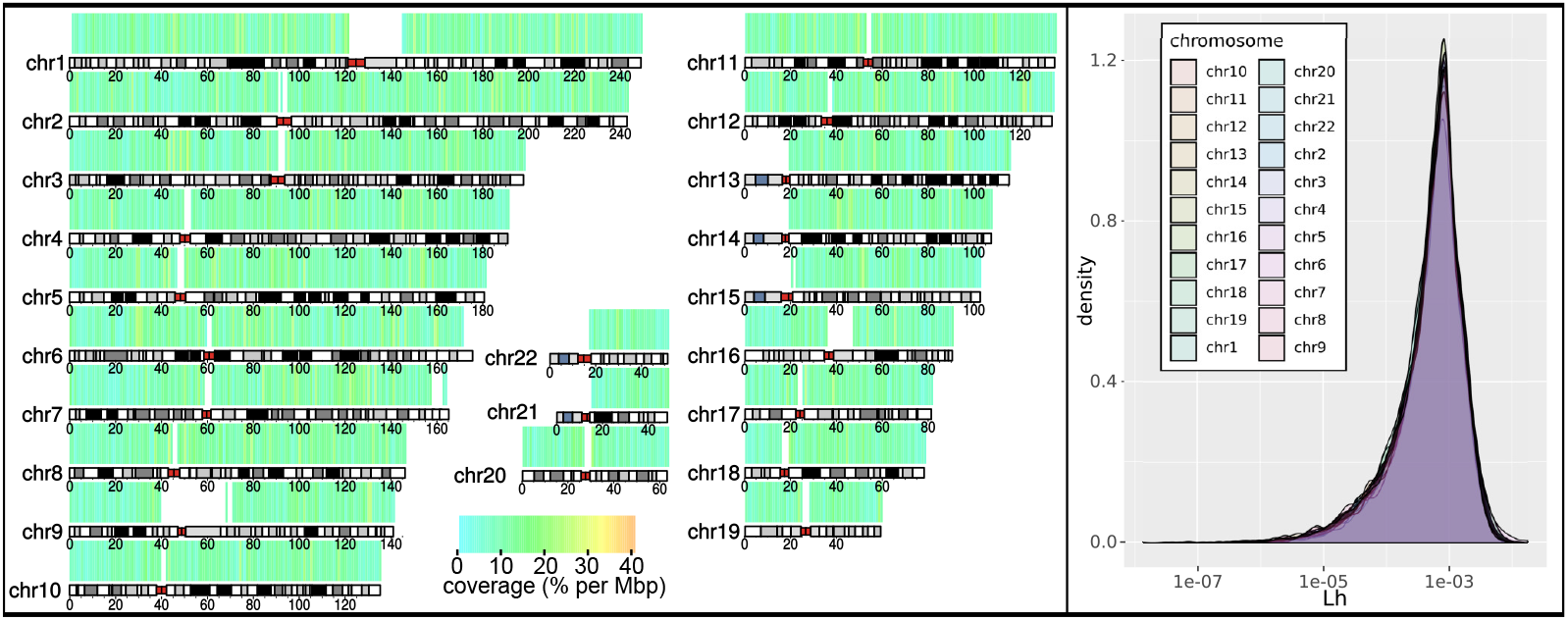
(Left) Genome coverage of haplotype blocks and (Right) Distribution of *L*[*h*], the length of haplotype blocks in cM. (Left) Genome coverage of haplotype blocks, in percent per Mbp. For each window of 1 Mbp, we computed the total length in bp covered by the 119,912 haplotype blocks (see section 2.2 for details).(Right) Distributions of *L*[*h*] (block length in cM), as defined by the genetic distance between bordering markers of each block. Blocks were defined over the 22 autosomal chromosomes using *δ* = 0.001*cM*, a parameter that account for the maximal length in cM between two consecutive markers in a same haplotype (see 2.2 and [21] for details). The distribution of block lengths is homogeneous across the 22 autosomal chromosomes.

### Haplotype tests exhibit different sensitivity compared to single SNP-based associations

We carried out a comparative study of the discovery power for the three haplotype tests. To do so, all significant phenotype-haplotype associations for the three tests were matched with the conventional SNP association results. In Fig. 3 top panel, the hits obtained with the three tests were matched with genotyped SNPs and with the imputed SNPs in the bottom panel of Fig. 3. For each significant phenotype-haplotype association, we used the lowest p-value of single-SNP associations observed in the SNPs of that haplotype block. For the complete-test and the single-test, if more than one haplotype per block was significantly associated, we selected the one with the lowest p-value. When comparing results among the three tests, we found that:

i. the block-test, that combines all haplotypes present in one block, leads to the lowest p-values for the strongest associations. However it appears globally more conservative than the single-SNP approach, particularly for associations in a range close to the genomic threshold (10^-8^ to 10^-10^). The negative intercept of the fitted linear trend (blue lines in Fig. 3) indicates that, in the significant blocks, p-values from this test are on average less significant than their single-SNP counterparts.
ii. the complete-test appears to be the most sensitive, with p-values on average being 3-fold lower than their single-SNP counterparts.
iii. the single-test performs similarly to the single-SNP approach, with p-values being slightly closer to those obtained with genotyped SNPs (*R*^2^ = 0.952) compared to those obtained with imputed SNPs (*R*^2^ = 0.944). Indeed, a large number of haplotypes are differentiated from the other haplotypes by a single SNP in the same block, and thus are expected to carry the same signal.

**Figure 3:**
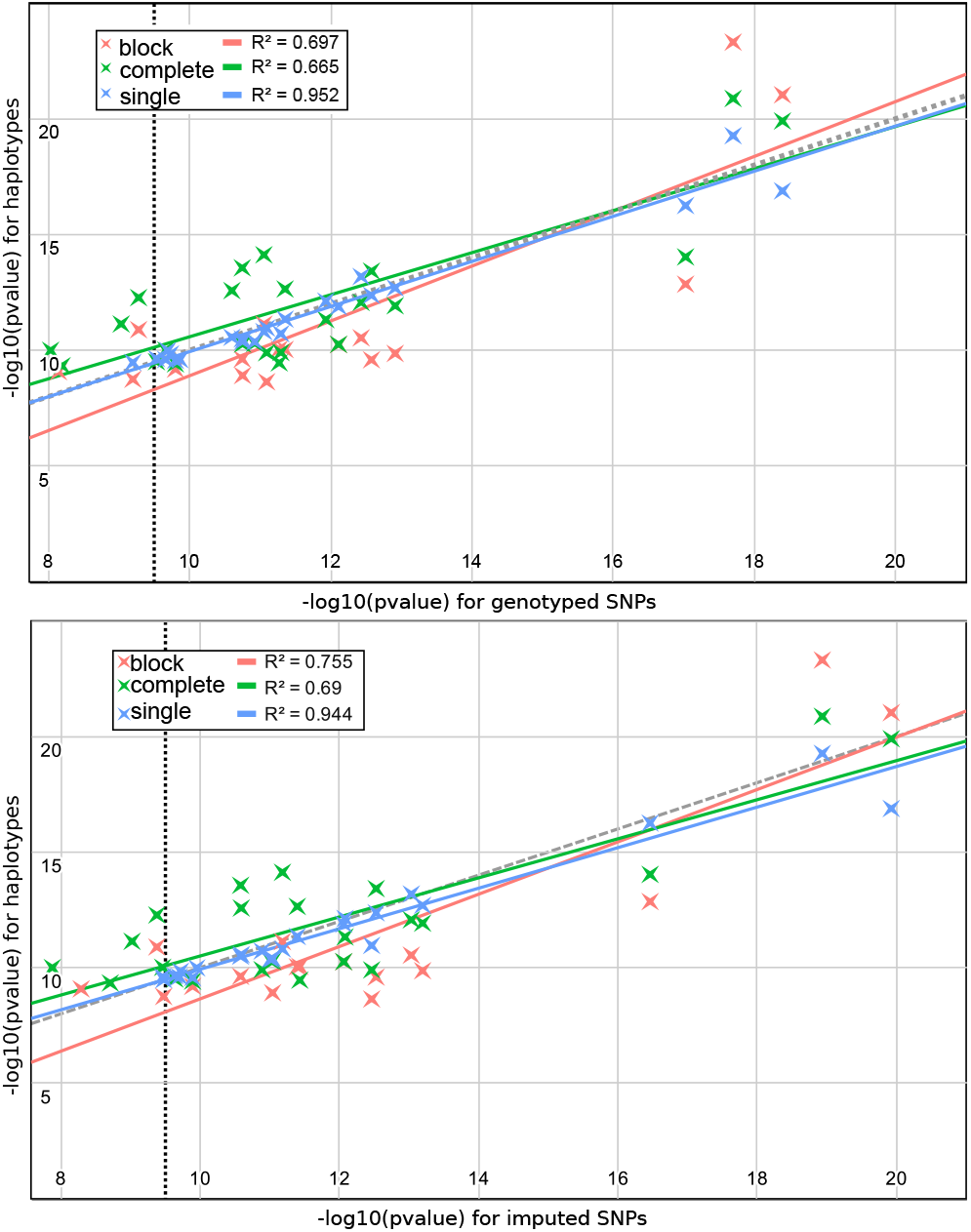
Comparison of p-values between main hits detected with haplotype association tests and their single-SNP approach counterpart. Comparison for main hits using the three proposed haplotype association tests (Y-axis) and single-SNP association (PLINK, X-Axis) with: (top) genotyped SNPs and (bottom) imputed SNPs. For each significant haplotype-phenotype association (Y-axis), the lowest p-value observed for the SNPs inside the block was reported (X-axis). For points above the grey line at x=y, p-values for haplotype associations test are lower than their single-SNPs counterpart. Blue lines indicates the fit of haplotype block test p-values (*y* = 1.13x – 2.67) with imputed SNPs and (*y* = 1.18*x* – 2.94) with genotyped SNPs; Red lines indicates the fit of complete model haplotype test p-values (*y* = 0.845*x* + 2.05) with imputed SNPs and (*y* = 0.908*x* – 1.5) with genotyped SNPs; Yellow lines indicates the fit for single haplotype test p-values (*y* = 0.877*x* + 1.16) with imputed SNPs and (*y* = 0.974*x* + 0.202) with genotyped SNPs

In Table 1 we used three available SNP datasets: (a; genotyped SNPs): SNPs called from the UK-Biobank arrays; (b; Imputed SNPs): imputed SNPs with a MAF>0.01 see [6]; (c; Rare SNPs): imputed SNPs with MAF<0.01. Similarly to Fig. 3, in order to match a given haplotype hit with a single-SNP hit in Table 1, we used the lowest p-value of single-SNP associations observed in the SNPs of that significant haplotype block.

**Table 1:**
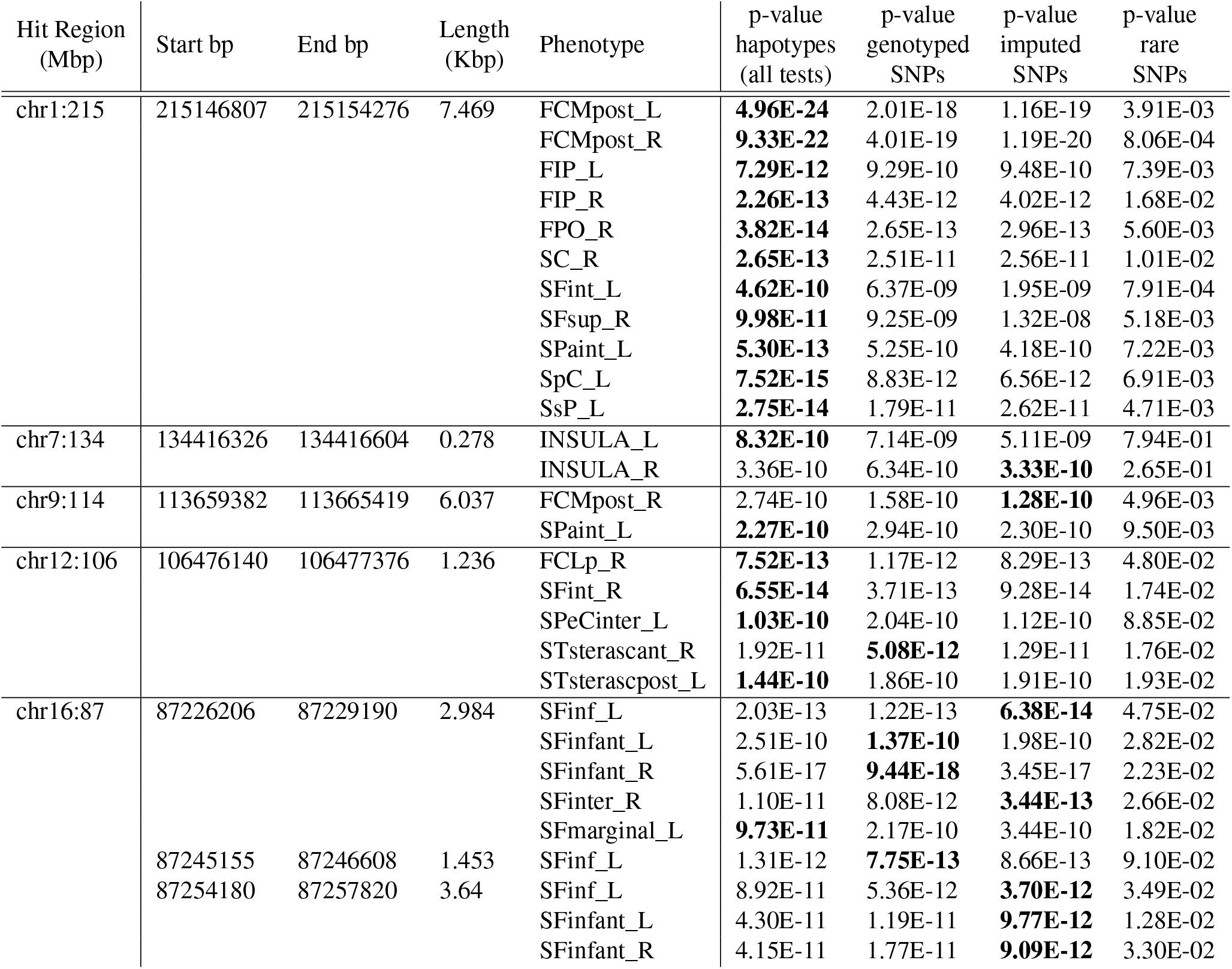
Significant hits found in haplotype blocks. For each block, the lowest p-value over the 3 haplotype tests is compared with the lowest p-value in the block for genotyped SNPs, imputed common SNPs (MAF > 1%) and rare SNPs (MAF < 1%). For each row, the lowest p-value is highlighted in bold. All positions are given for GRCh37/hg19 assembly.

| Hit Region<br>(Mbp) | Start bp | End bp | Length<br>(Kbp) | Phenotype | p-value<br>haplotypes<br>(all tests) | p-value<br>genotyped<br>SNPs | p-value<br>imputed<br>SNPs | p-value<br>rare<br>SNPs |
| --- | --- | --- | --- | --- | --- | --- | --- | --- |
| chr1:215 | 215146807 | 215154276 | 7.469 | FCMpost_L | **4.96E-24** | 2.01E-18 | 1.16E-19 | 3.91E-03 |
|  |  |  |  | FCMpost_R | **9.33E-22** | 4.01E-19 | 1.19E-20 | 8.06E-04 |
|  |  |  |  | FIP_L | **7.29E-12** | 9.29E-10 | 9.48E-10 | 7.39E-03 |
|  |  |  |  | FIP_R | **2.26E-13** | 4.43E-12 | 4.02E-12 | 1.68E-02 |
|  |  |  |  | FPO_R | **3.82E-14** | 2.65E-13 | 2.96E-13 | 5.60E-03 |
|  |  |  |  | SC_R | **2.65E-13** | 2.51E-11 | 2.56E-11 | 1.01E-02 |
|  |  |  |  | SFint_L | **4.62E-10** | 6.37E-09 | 1.95E-09 | 7.91E-04 |
|  |  |  |  | SFsup_R | **9.98E-11** | 9.25E-09 | 1.32E-08 | 5.18E-03 |
|  |  |  |  | SPaint_L | **5.30E-13** | 5.25E-10 | 4.18E-10 | 7.22E-03 |
|  |  |  |  | SpC_L | **7.52E-15** | 8.83E-12 | 6.56E-12 | 6.91E-03 |
|  |  |  |  | SsP_L | **2.75E-14** | 1.79E-11 | 2.62E-11 | 4.71E-03 |
| chr7:134 | 134416326 | 134416604 | 0.278 | INSULA_L | **8.32E-10** | 7.14E-09 | 5.11E-09 | 7.94E-01 |
|  |  |  |  | INSULA_R | 3.36E-10 | 6.34E-10 | **3.33E-10** | 2.65E-01 |
| chr9:114 | 113659382 | 113665419 | 6.037 | FCMpost_R | 2.74E-10 | 1.58E-10 | **1.28E-10** | 4.96E-03 |
|  |  |  |  | SPaint_L | **2.27E-10** | 2.94E-10 | 2.30E-10 | 9.50E-03 |
| chr12:106 | 106476140 | 106477376 | 1.236 | FCLp_R | **7.52E-13** | 1.17E-12 | 8.29E-13 | 4.80E-02 |
|  |  |  |  | SFint_R | **6.55E-14** | 3.71E-13 | 9.28E-14 | 1.74E-02 |
|  |  |  |  | SPeCinter_L | **1.03E-10** | 2.04E-10 | 1.12E-10 | 8.85E-02 |
|  |  |  |  | STsterascant_R | 1.92E-11 | **5.08E-12** | 1.29E-11 | 1.76E-02 |
|  |  |  |  | STsterascpost_L | **1.44E-10** | 1.86E-10 | 1.91E-10 | 1.93E-02 |
| chr16:87 | 87226206 | 87229190 | 2.984 | SFinf_L | 2.03E-13 | 1.22E-13 | **6.38E-14** | 4.75E-02 |
|  |  |  |  | SFinfant_L | 2.51E-10 | **1.37E-10** | 1.98E-10 | 2.82E-02 |
|  |  |  |  | SFinfant_R | 5.61E-17 | **9.44E-18** | 3.45E-17 | 2.23E-02 |
|  |  |  |  | SFinter_R | 1.10E-11 | 8.08E-12 | **3.44E-13** | 2.66E-02 |
|  |  |  |  | SFmarginal_L | **9.73E-11** | 2.17E-10 | 3.44E-10 | 1.82E-02 |
|  | 87245155 | 87246608 | 1.453 | SFinf_L | 1.31E-12 | **7.75E-13** | 8.66E-13 | 9.10E-02 |
|  | 87254180 | 87257820 | 3.64 | SFinf_L | 8.92E-11 | 5.36E-12 | **3.70E-12** | 3.49E-02 |
|  |  |  |  | SFinfant_L | 4.30E-11 | 1.19E-11 | **9.77E-12** | 1.28E-02 |
|  |  |  |  | SFinfant_R | 4.15E-11 | 1.77E-11 | **9.09E-12** | 3.30E-02 |

The comparative analysis given in Table 1 revealed the following salient points:

i. Haplotype association tests have similar or better detection power than imputed or genotyped SNP counterparts. Indeed, in the majority of hits, there was at least one haplotype model that was more significant than the single-SNP test (18 vs. 11). Moreover, for 9 phenotype/genotype significant associations, the haplotype p-values were at least 10-fold lower than their single-SNP counterparts while the opposite was true only once.
ii. For strong associations involving several haplotypes in the same block, haplotype approaches can lead to p-values that are several order lower, for example, haplotypes association in block chr1:215146807-215154276 with FCMpost Left leads to p-values in the range of 10^-24^ compared to 10^-19^ with single-SNP associations. Additionally, for 6 other phenotypes, p-values obtained using haplotype associations are also 2 or 3 orders of magnitude lower.
iii. Imputed rare variants do not exhibit any significant associations. Indeed, the smallest p-values associated with rare variants are 6 to 20 orders of magnitude higher than the significant haplotype hits.

### False Positive Rates in the Genome-wide haplotype association tests

Table 2 shows an estimation, for each test, of *FPR_T_* = *N_FP_/N_t_*, the FPR for the null distributions that preserve the correlation among phenotypes and within the genome (third scenario described in Section 2.3.4). The Q-Q plots presented in Fig. 4 indicate that in this permutation scenario, we do not observe an inflation of p-values under the null hypothesis. When using Bonferroni correction for significance thresholds (see 2.3.4), none of the three tests seem to show a systematic inflation of FPR (see Table 2). In Section Supplementary Material 5, we investigated FPR in the two other scenarios of null hypotheses as described in Section 2.3.4 and reported results in Fig 3 (for scenario 1) and Table 1 (for scenario 2). We show that in the second permutation scenario, out of the two phenotypes (SPoCsup_left and SRh_left) showing systematic inflated FPR, none of them were involved in significant hits on real data. In Fig 3, we did not observed any inflation of FPR under the first permutation scenario. In addition, we did not observed any False Positive.

**Figure 4:**
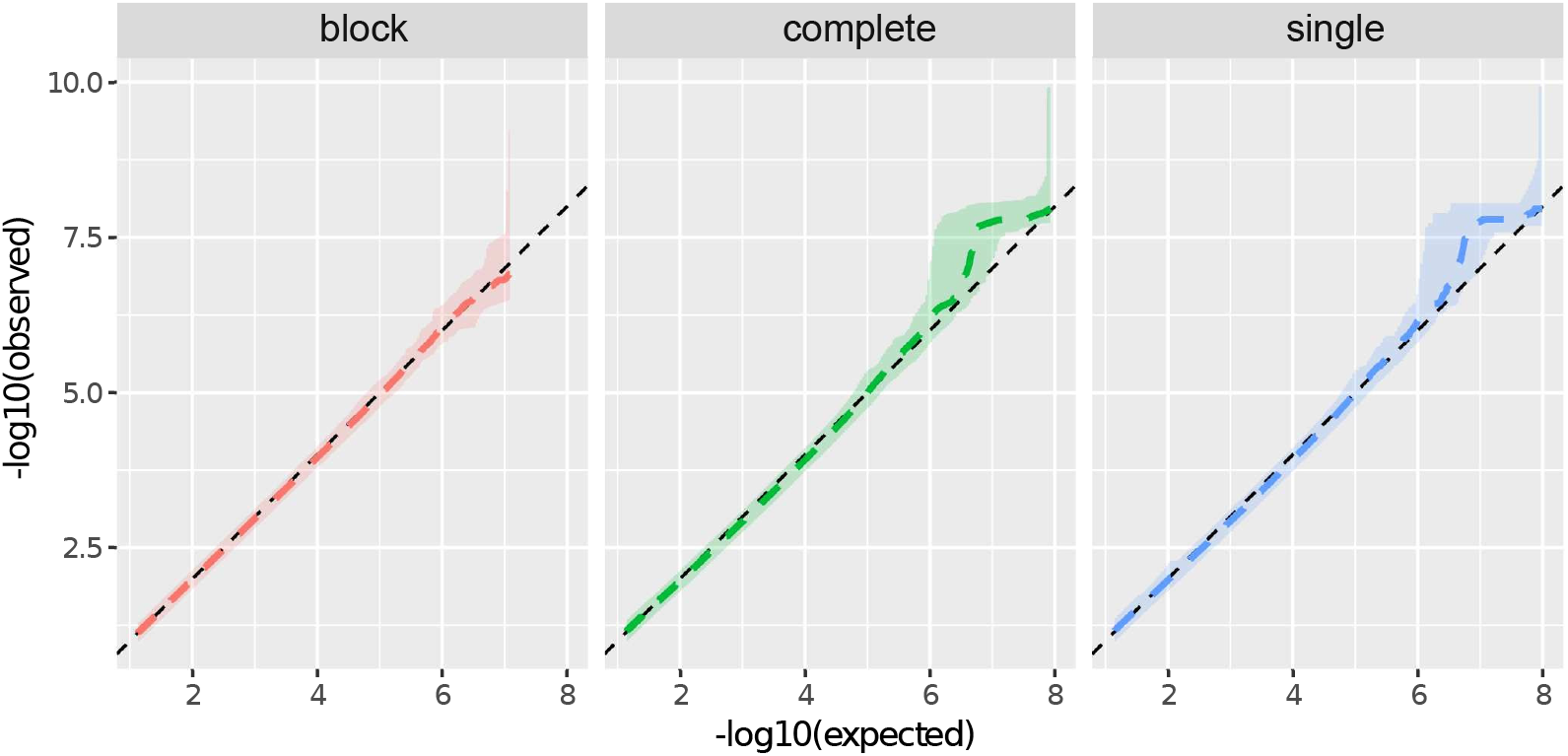
Aggregation of QQ-plots for 10 runs of permutated datasets. Aggregation of QQ-plots for 10 runs of permutated datasets (see 2.3.4, third scenario). From left to right: block-test, complete-test and single-test. The shaded area define the hull (minimum and maximum values) of the Q-Q plots for the 10 runs and the average Q-Q plot is given by the coloured, dotted line. The black, dashed line at *y* = *x* indicates where the expected distribution lies.

**Table 2:** False Positive Rate under null hypothesis for 10 runs of permutations using the third scenario (see section 2.3.4). For each run of permutation, tests were corrected by the Bonferroni procedure *α′* = *α/N_T_* for *N_T_* tests with risk *α* = 0.05 i.e. the FPR should be lower than 1/*N_T_* in 5% of the runs.

| Test | Average<br>number<br>of tests<br>per run | Average<br>discovery<br>threshold<br>$\alpha' = \alpha/N_T$ | Average<br>FPR | Number of<br>run with<br>$\text{FPR} \geq 1/N_T$ | Min FPR | Max FPR |
| --- | --- | --- | --- | --- | --- | --- |
| Block | 14,673,826 | 3.4E-09 | 6.8E-9 | 1 | 0 | 6.8E-8 |
| Complete | 107,683,292 | 4.6E-10 | 1.8E-9 | 1 | 0 | 9.2E-09 |
| Single | 124,951,506 | 4.0E-10 | 1.6E-9 | 1 | 0 | 8.0E-09 |

## 4 Discussion

### Box-Cox transformation normalizes phenotypes across brain

We used Box-Cox transformation of sulcal opening values to ensure a spatially homogeneous distribution of the opening measurements across the brain. The spatial distribution of the λ parameter across the brain highlights the necessity of such a transformation. This procedure ensures a homogeneous specificity of the analysis across the brain sulci. Indeed, the spatial pattern in the λ values does not reflect the spatial distribution of the FPR. In the analysis on permuted datasets, we observed an overall homogeneous behaviour across the brain for the three haplotype tests (see Section 1).

### Choosing δ to define blocks

We set δ parameter to the most conservative values among the ones we studied. The objective is to ensure the reliability of the phase along the block. We observed that the distribution of the block genetic length *L*[*h*] for this value is the most homogeneous across the different chromosomes. Section Supplementary Material 4 shows that with δ increasing from 0.001 to 0.025, the distribution becomes bi-modal and less homogeneous across the chromosomes. With higher values of *δ*, *L*[*h*] exhibits heterogeneous distributions depending on the chromosome. Moreover, with a higher δ value, a larger number of recombination is allowed within the same haplotype block which may lead to a lower sensitivity.

### Sulcal opening patterns are associated with haplotypes

In this study we show significant associations of sulcal opening measurements with haplotypes in a genome-wide approach; bringing about genome wide inferences that differ from a local haplotype fine-mapping operation performed after a classical genome wide single-SNP analysis. Along with the previous findings in the upstream region of KCNK2 (chr1:215MB) [10], the haplotype block test revealed 4 other hits on chromosomes 7, 9, 12 and 16, detailed in Table 3. For the following genes, located in these regions, that are predominantly expressed in the brain: LPAR1 (chr9); NUAK1 (chr12), lrcRNA RP11-178L8.8 and FBXO31 (chr16), we provide an overview of expression data, publicly available in Supplementary Material 6. We also display brain regions associated with each hit in Supplementary Material 7. Except for chr9:114, all hits reported on Table 1 are associated bilaterally with sulci such as (FCMpost_L, FCMpost_R) for chr1:215, (STsterascant_R, STsterascpost_L) for chr12:106 and (SFinfant_L, SFinfant_R) for chr16:87.

**Table 3:** Genes or regions of interest related to haplotypes significantly associated with the opening of at least one sulcus. All positions are given for GRCh37/hg19 assembly.

| Haplotype position | Gene / Region position | Description |
| --- | --- | --- |
| chr1:<br>215146807-215154276 | upstream of<br>KCNK2 | eQTL of KCNK2 [10] |
| chr7:<br>13441632-134416604 | chr7:<br>134346283-134446935<br>chr7:<br>129202420-138543686 | autism susceptibility region<br>7q32.3-q33 [24]<br>AUTS1 region :<br>D7S530 to D7S684 [25, 26] |
| chr9:<br>113659382-113665419 | chr9:<br>113635543-113800981<br>(intronic) | LPAR1 : Homo sapiens lysophosphatidic acid receptor 1, transcript variant 2, mRNA. Protein, most expressed in Brain - Spinal cord (cervical c-1) (GTEx V7) |
| chr12:<br>106476140-106477376 | chr12:<br>106457118-106533811<br>(intron,<br>exon 3 & 4) | NUAK1 : Homo sapiens NUAK family, SNF1-like kinase, 1, mRNA. Protein, most expressed in Brain - Frontal Cortex (BA9) (GTEx V7), involved in axon branching [31] |
| chr16:<br>87226206-87257820 | chr16:<br>87360593-87361190<br>(102kpb,<br>upstream)<br>chr16:<br>87360593-87422364<br>(102kpb,<br>downstream) | RP11-178L8.8 (AC010531.7): novel transcript, antisense to FBXO31 and C16orf95 readthrough LncRNA, most expressed in Brain - Cerebellum(GTEx V6)<br>FBXO31 : F-box protein 31,mRNA. Protein, most expressed in Brain - Cerebellum(GTEx V8) |

Among the bilateral associations, the hit chr7:134 stands out: its associated brain areas - the insula (left and right) - are not associated with any other genome region. Further studies could be motivated to investigate this genomic region since the insula is a cortical region known to integrates emotional, cognitive, and motivational signals, and that haplotypes of the block chr7:13441632-134416604 are located in a region carrying several genes and markers associated with Autism Spectrum disorders ([24, 25, 26], see details in Table 3).

### Multiple testing inside a block

For complete-test and single-test, the number of independent tests probably lies between the number of blocks and the number of haplotypes. For example, a block could contain a main, very common haplotype, and several rare haplotypes associated with low-frequency SNPs. In such a case, it is likely that the association p-values of the different haplotypes within the same block will be dependent. The bump in the Q-Q plots in Fig. 4 (middle and right panels) could come from the sample with the main population carrying the main haplotype, and only a few individuals driving an association for the minor haplotypes. In a similar situation where many haplotypes exhibit the same structure in the population, Howard et al. [27] proposed a correction using an estimation of the number of independent haplotypes, which would be less stringent than the Bonferroni correction.

### Disentangling multiple neighbouring single SNP hits using haplotypes

Out of the three haplotype tests studied, the complete model haplotype test turned out to be the most sensitive overall. However, results show that in the presence of a complex LD background, like in the region upstream of KCNK2, the haplotype modelling underlying the block-test model produces p-values several orders of magnitude lower. This result calls for the development of innovative multivariate approaches based on haplotypes, such as those the authors have started to propose [28].

### Limitations of the study

We choose to use a low *δ* value in order to obtain the most reliable haplotypes. One drawback is that this also allows higher LD between the blocks, meaning a greater correlation between blocks than with higher *δ* values. Using Bonferroni correction for multiple testing is expected to be conservative in the presence of such dependencies.

In our study of FPR, we purposely focus on realistic null distributions of measurements, that exhibit the true correlation *between all the phenotypes and along the whole genome*. To fully control the FPR, one would need to run a larger number of runs which is computationally intensive and out of reach using our realistic null distributions. However, with the number of permutations used, we could reasonably detect when a large inflation of FPR occurs.

By comparing with single-SNPs analyses, we did not directly estimate the discovery power of our tests. To our knowledge, for whole-genome haplotype associations, there is no practical way to create a synthetic and realistic ground truth signal for multiple correlated phenotypes.

### Conclusion and future works

In the context of imaging-genetics, we studied three haplotype tests to find associations between haplotypes and quantitative traits measured on brain MRI. Beyond the process of extracting features from the imaging-genetics data, we normalized the measurement distributions of the sulcal openings using Box-Cox transformations and obtained spatially homogeneous distributions across the brain. We compared three haplotype association tests and achieved the best performance with the haplotype block model test (lowest p-values observed) and the complete model haplotype test (best sensitivity). These two tests use a multivariate model of haplotypes that account for the whole set of haplotypes present in a given block. We show that these tests outperformed the single-SNP based classical approach for haplotype blocks with association hits close to the genomic threshold (10^-8^ to 10^-10^), specifically in the case of haplotype blocks that exhibit complex LD structures. Based on the results of this work, we do not argue for the systematic use of haplotype modelling over GWAS based on imputed variants. More particularly, in the case of the UK Biobank, all samples come from a homogeneous population with an extensive imputation panel [6]. In this case, the discovery power of haplotypes is often in the same range as that of imputed SNPs. However, in the context of imaging-genetics, we show that using haplotypes showed more reliable results: significant haplotypes were associated with a larger number of phenotypes, increasing the explanatory value of each discovery. This study relies on the definition of blocks using a genetic map and a single value of δ. Future works could take a step further and locally define block boundaries in order to find the more relevant ones in terms of association [29, 30].

## Acknowledgment

This work was funded by CEA - Université Paris-Saclay and FRM grant number DIC20161236445

## Conflict of interest

Authors declare no conflict of interest.

## Supplementary Material 1 Box-Cox transformation of Sulcal Opening

**Supplementary Figure 1:**
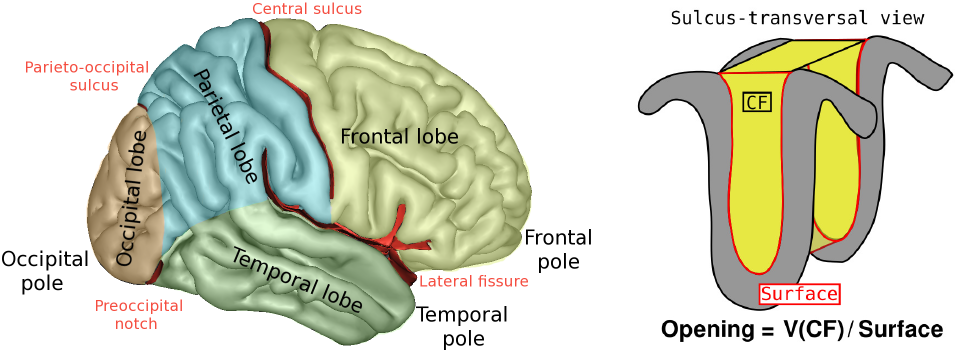
(left) Main brain structures. Sulci are the main furrows. Here only two sulci are labelled. For example, Central Sulcus is separating Frontal and Parietal lobes (Blausen Medical Encyclopedia). (right) Sulcus opening, a measure of the average sulcus width, is computed as the ratio of the volume of Cerebrospinal Fluid to the surface of the sulcus.

For each subject, 123 labelled sulci were extracted from T1-weighted images. For each sulcus, a measure of its width - a feature called opening - is computed as the ratio of the volume of the Cerebrospinal Fluid the sulcus contains to its surface. For each sulcus, after adjusting for age and sex using linear regression, we identified and excluded outliers in the residual distribution using the robust interquartile range (IQR) method [17]. The distributions of sulcal opening values could exhibit deviation from the normal distribution. We evaluated these deviations and normalized them using a one-parameter Box-Cox transformation (power transformation): Let **Y** and 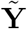 be the vectors of the initial and Box-Cox transformed sulcal opening values respectively. The one-parameter Box-Cox transformation is the following:

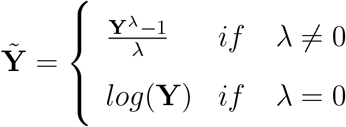

## Supplementary Material 2 Haplotype count matrix

The following shows an example of Haplotype count matrix **H**, obtained with 3 phased SNPs and 3 subjects: the matrix **H** accounts for the number of copies, per subject, of the two observed alternative haplotypes *h*_1_ = [101] and *h*_2_ = [010]. In our example, subject *S*_1_ is homozygous for haplotype *h*_0_ and subjects *S*_2_ and *S*_3_ are heterozygous with haplotypes *h*_1_ and *h*_2_, and haplotypes *h*_0_ and *h*_2_ respectively.

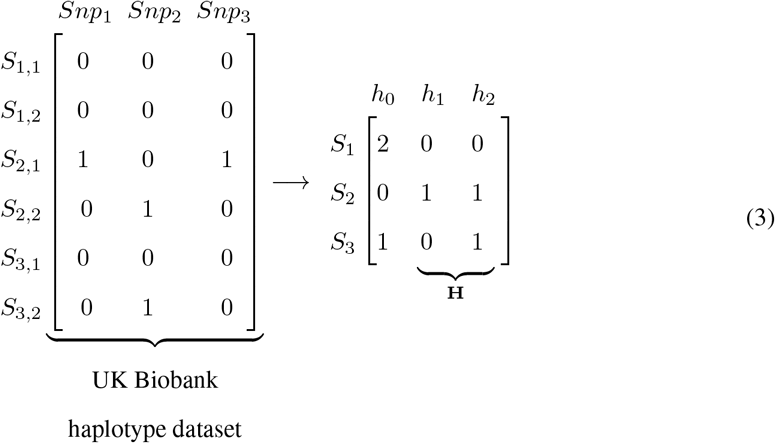

## Supplementary Material 3 Univariate single-SNP test classically used in GWAS

The test for each of the *p* SNPs in classical GWAS for quantitative traits uses the following linear model:

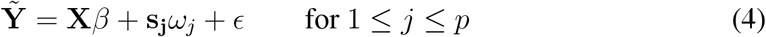

with 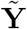 the phenotype vector, **X** the matrix of covariates and **s_j_** the count vector of minor alleles for the SNP *s_j_*, included with fixed effects *β* and *ω_j_* respectively, and *ϵ* the error vector. Using PLINK, we computed a standard two-sided *p*-value of the *t*-statistic to test the following null hypothesis:

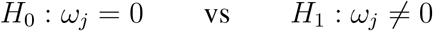

## Supplementary Material 4 Length of haplotype blocks along the genetic map

**Supplementary Figure 2:**
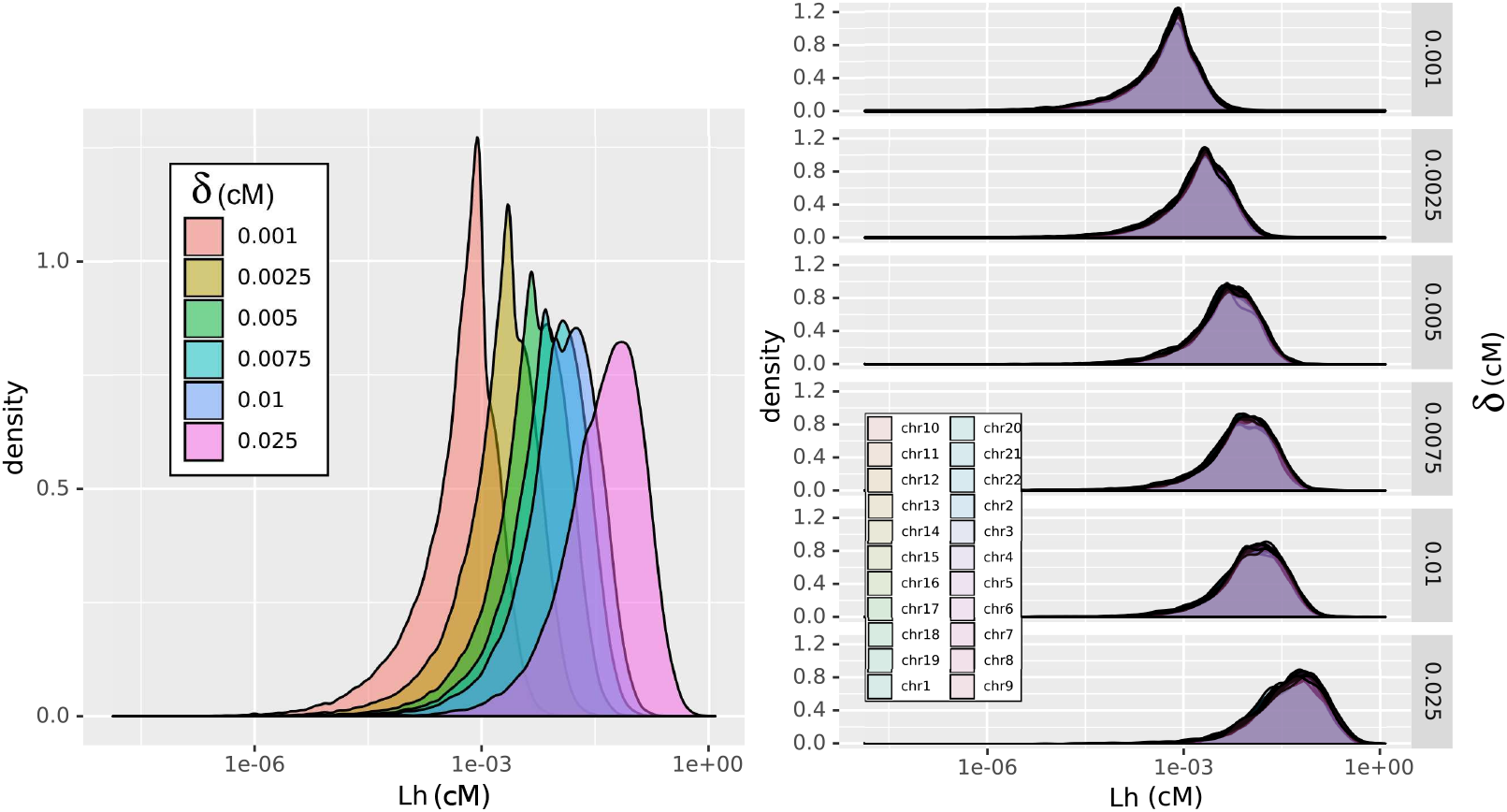
Distribution of haplotype block length, in cM for increasing value of parameter *δ*

## Supplementary Material 5 False Positive Rate under null hypothesis

**Supplementary Figure 3:**
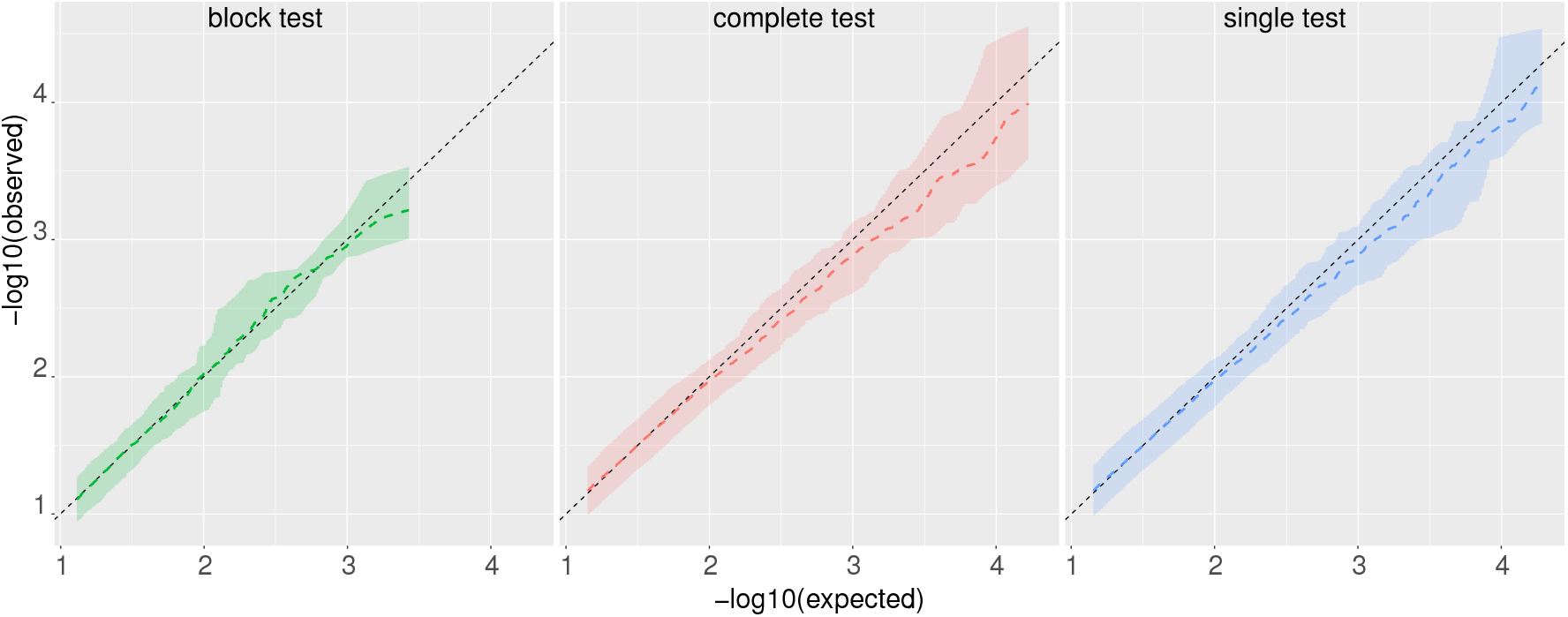
Aggregated QQ-plots under the null-hypothesis while preserving the correlation within the haplotype blocks (see section 2.3.4 first scenario, for details) using block-test, complete-test and single-test. We selected three phenotypes: two that represent the range of λ values (see 2.2 for details) and one for the most significant association found with the original dataset. For each test, we computed a QQ-plot for each of the three phenotypes. The shaded area define the hull (minimum and maximum values) of the Q-Q plots for the 3 phenotypes and the average Q-Q plot is given by the coloured, dotted line. The black, dashed line at *y* = *x* indicates where the expected distribution lies. For the three tests and the three phenotypes, the procedure did not produce any False Positive using *α* = 0.05 and Bonferroni correction.

**Supplementary Table 1:**
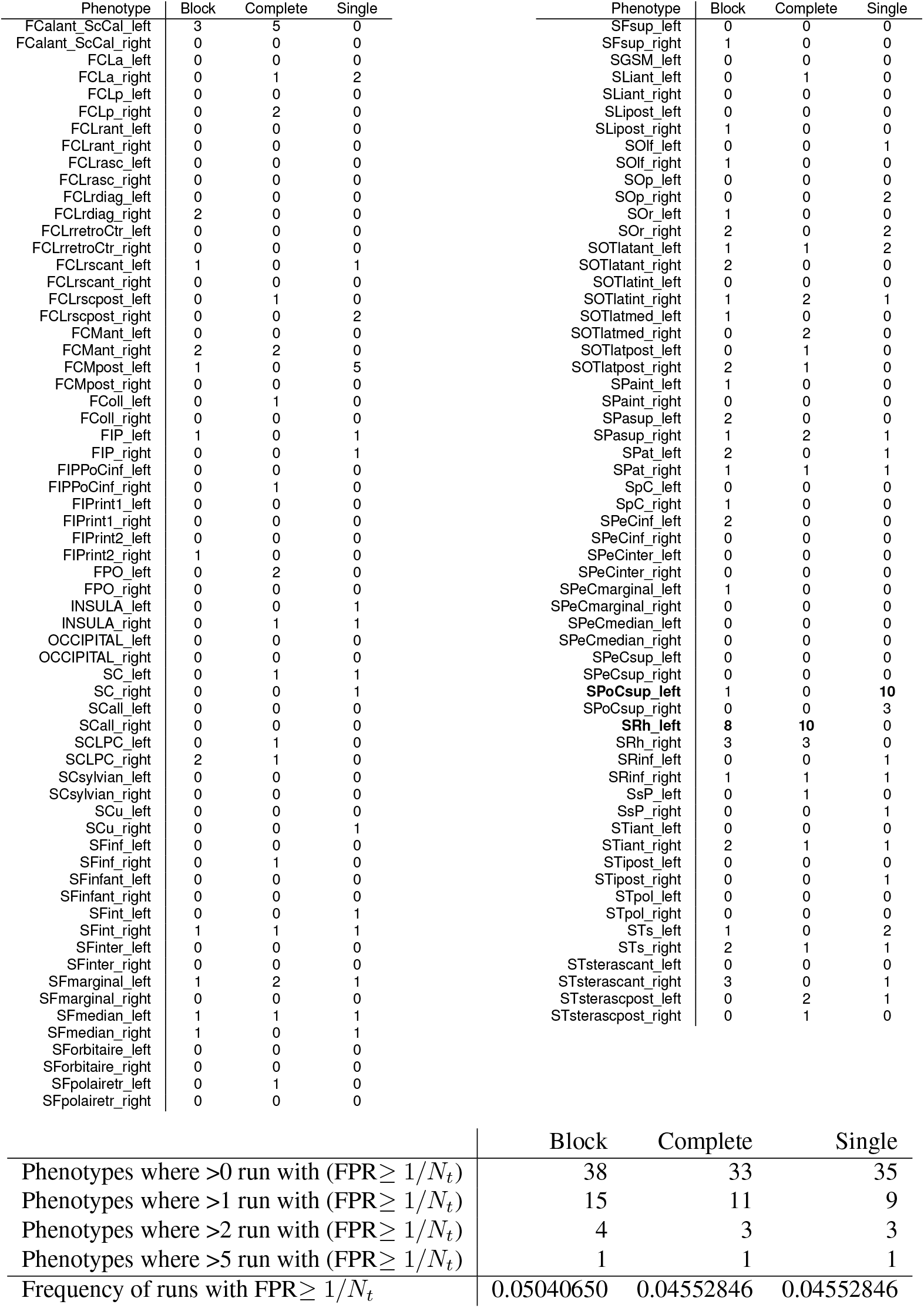
Study of the False Positive Rate under the null hypothesis for 10 runs of permutation using the second scenario (see section 2.3.4 for details). P-values are corrected using Bonferroni (*α*/*N_t_*) for *N_t_* hypotheses, and discovery threshold set to *α* = 0.05. The top table shows the number of runs (out of 10) where we observed an estimated FPR ≥ 1/*N_t_*. The bottom table aggregates these counts per test, and shows that, when we pool the FPR results of all permuted phenotypes, we seem to control the family-wise error rate at 5% under the global null hypothesis for each of the three tests. Two phenotypes raise concern regarding inflation of FPR: SPoCsup_left and SRh_left but were not found significantly associated with any haplotype in the real dataset.

## Supplementary Material 6 Tissue expression of gene related to significant hits

The hit located on chr.7q is found only associated to Insula (left and right) openning. The insula is a cortical region known to integrates emotional, cognitive, and motivational signals. Haplotypes of the block chr7:13441632-134416604 are located in a genomic region comprising several genes and markers associated with Autism Spectrum disorders (see Table 3). This hit could carry a signal distinct from the others because it is the only single phenotype (bilateral) - single haplotype block significant association.

Haplotype found on chr9:113 are located within an intronic region of LPAR1 gene, which is encoding for a protein used in cell signalling, notably in inhibition of neuroblastoma cell differentiation. Stankoff *et al*. [32] showed that transcripts are not detected during early stages of oligodendroglial development, but are expressed only in mature oligodendrocytes, shortly before the onset of myelination. Transcript are expressed in different brain tissues in GTEx v8 (see Fig 4)

Haplotypes in block on chr.12:106476140-106477376 are located within the gene NUAK1, covering large intronic parts as well as 2 small exons. NUAK1 codes for a protein involved in cell proliferation, for example in arborization of mammalian neurons, regulating axon branching [31]. Particularly, it is expressed in several brain tissues (see Fig. 5).

Haplotypes found in the chr16:87Mbp region are associated with several sulci located on both sides of the frontal region, in particular the Inferior Frontal Sulcus (SFinf). Located within gene C16orf95, haplotypes are 100kbp upstream of RP11-178L8.8 (or AC010531.7), a non-coding lncRNA mostly expressed in the brain (GTEx v6). This locus is antisens to FXBO31, a gene predominently expressed in Brain tissues in GTEx v8 (see Fig. 6).

**Supplementary Figure 4:**
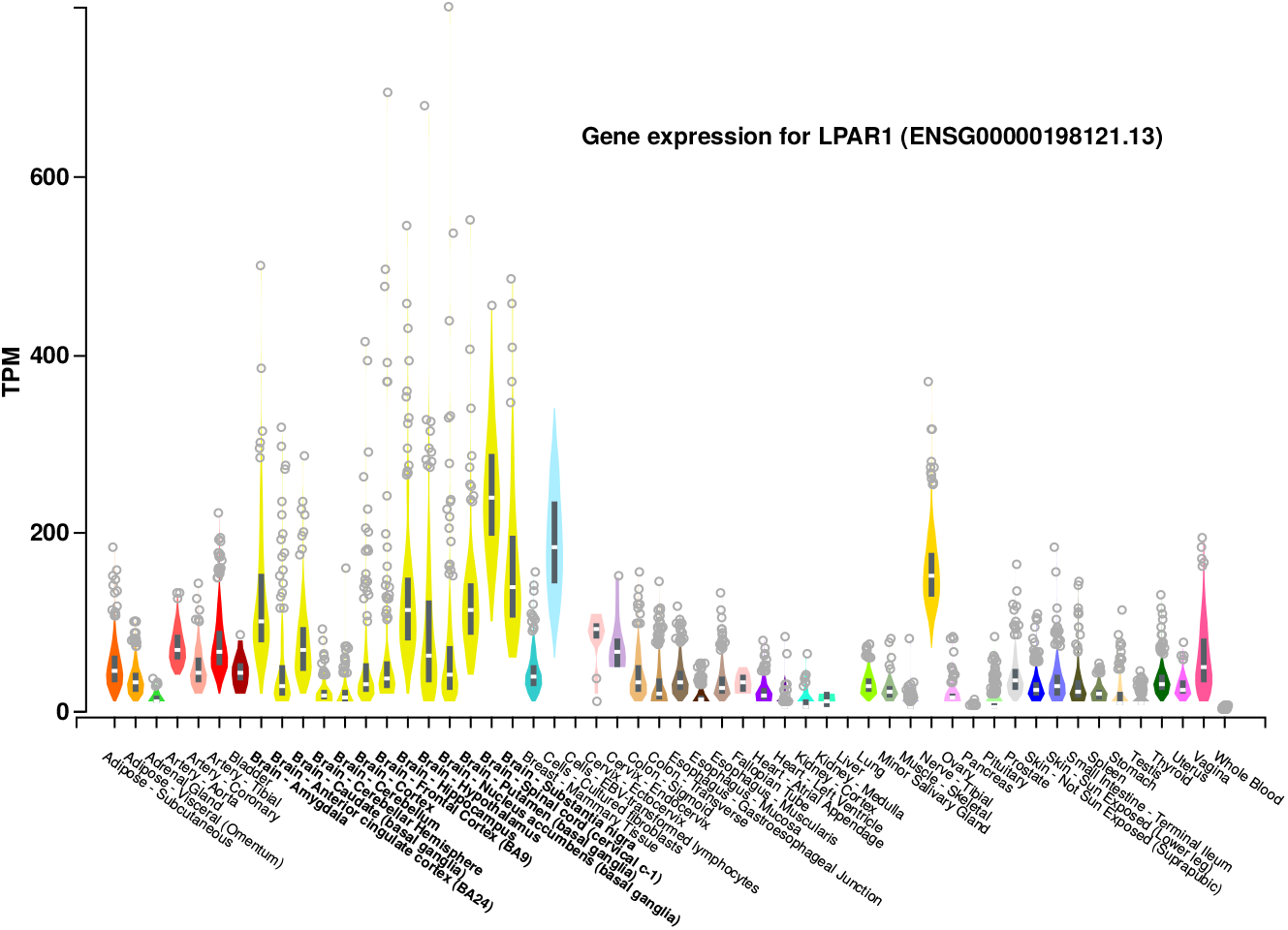
Expression of LPAR1 gene in different tissues of the brain (GTEx v8), related to top hit on chromosome 9.

**Supplementary Figure 5:**
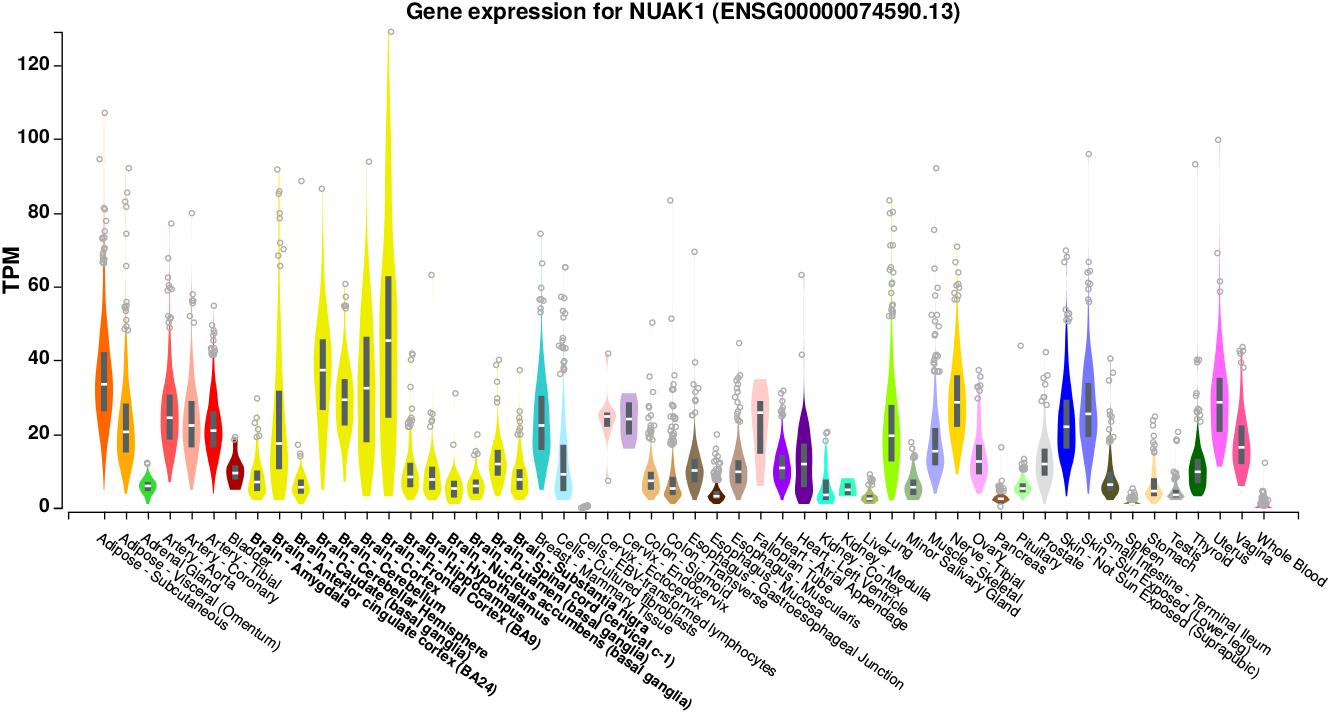
Expression of NUAK1gene in different tissues of the brain (GTEx v8), related to top hit for chromosome 12.

**Supplementary Figure 6:**
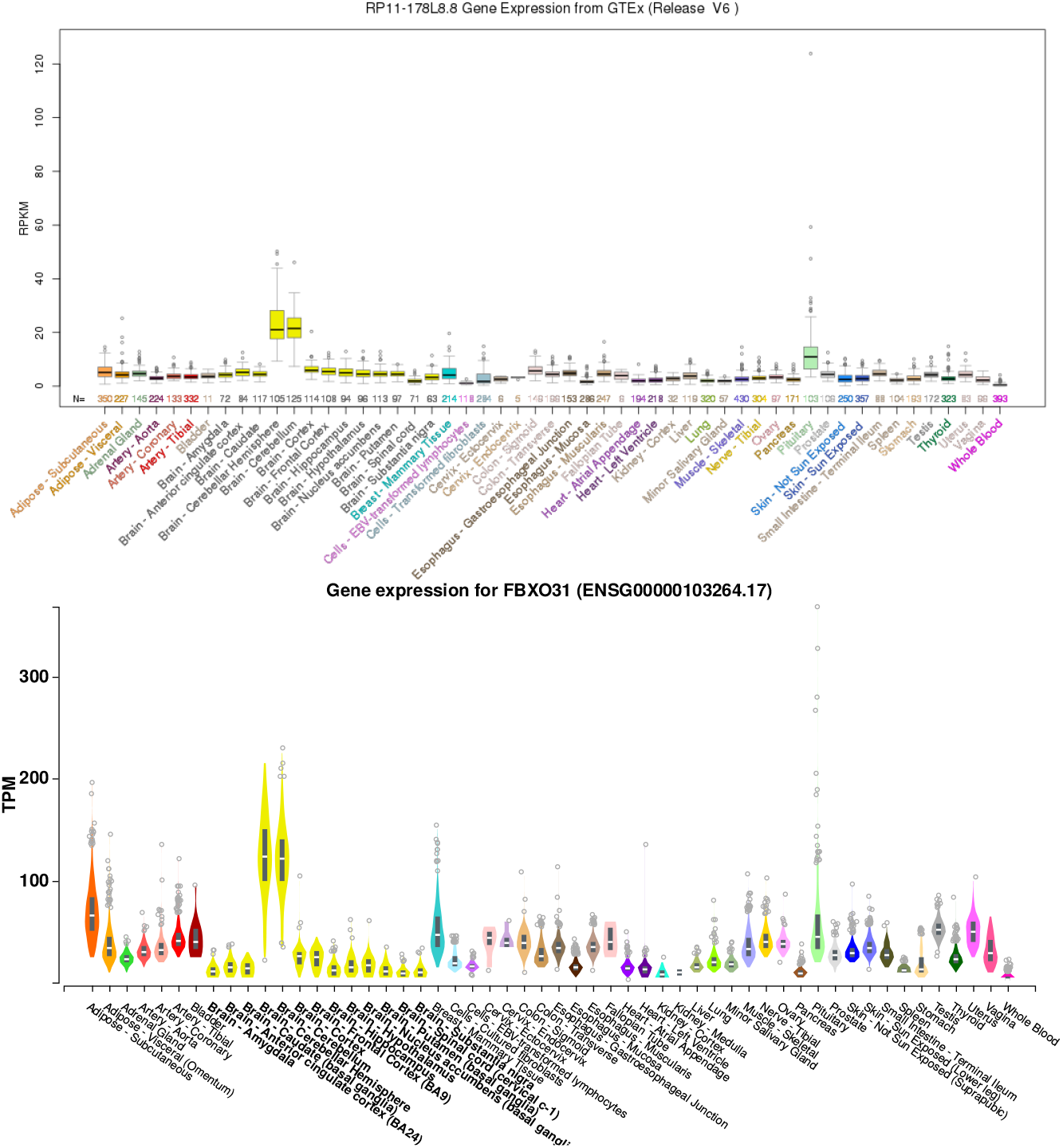
(Top) Expression of antisens lrcRNA RP11-178L8.8 in different tissues of the brain (GTEx v6), 102kbp downstream of top hit for chromosome 16 (Bottom) Expression of FBXO31 gene in different tissues of the brain (GTEx v8), 102kbp upstream of top hit for chromosome 16.

## Supplementary Material 7 Sulci of which openning is significantly associated with at least one genomic region

**Supplementary Figure 7:**
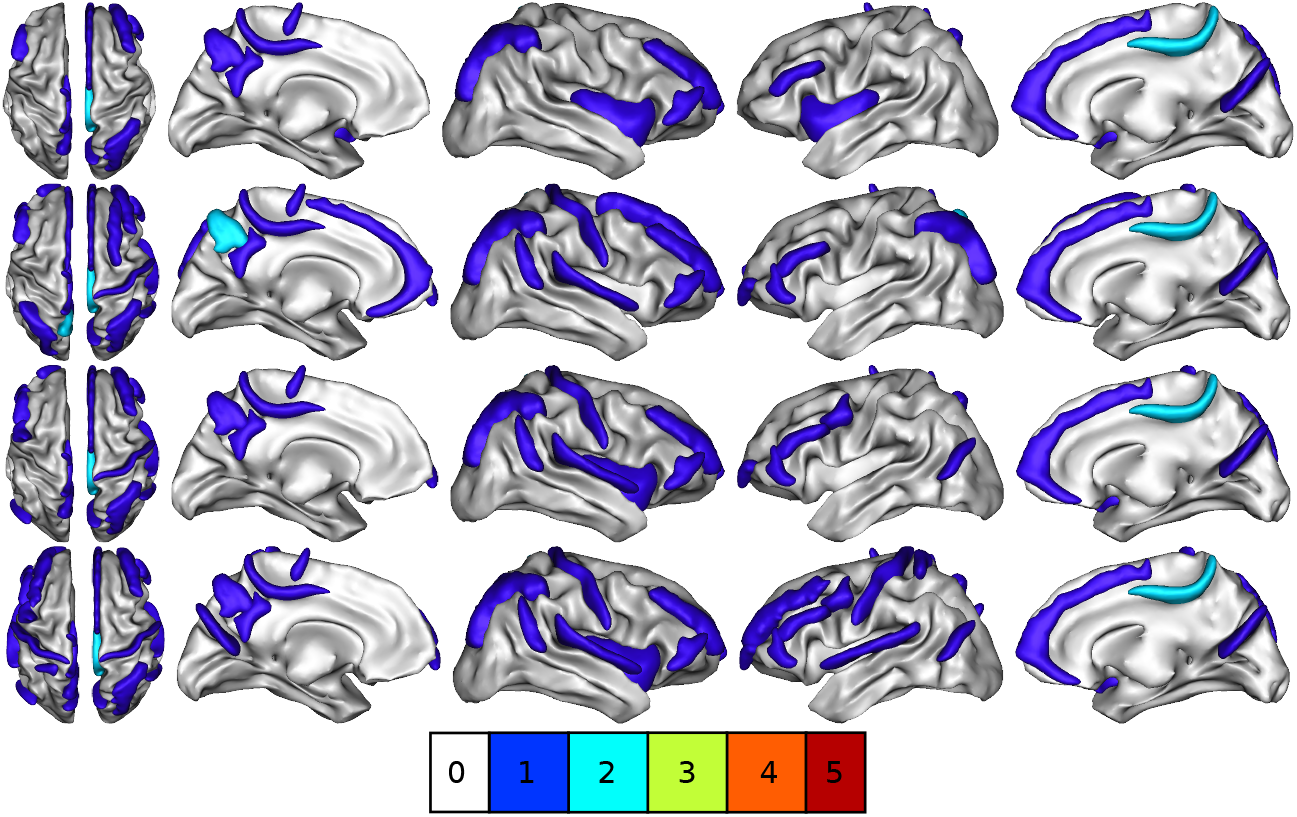
Brain sulci coloured according to the number of significant hits. Hits - genomic region of 1 Mbp with at least one significant p-value among the three tests. Sulci colored in dark blue color are associated with one hit, light blue are associated with 2 hits (chr1:215 and chr9:114). In rows, from top to bottom, hits are determined using block-test, complete-test, single-test and imputed SNPs with PLINK.

**Supplementary Figure 8:**
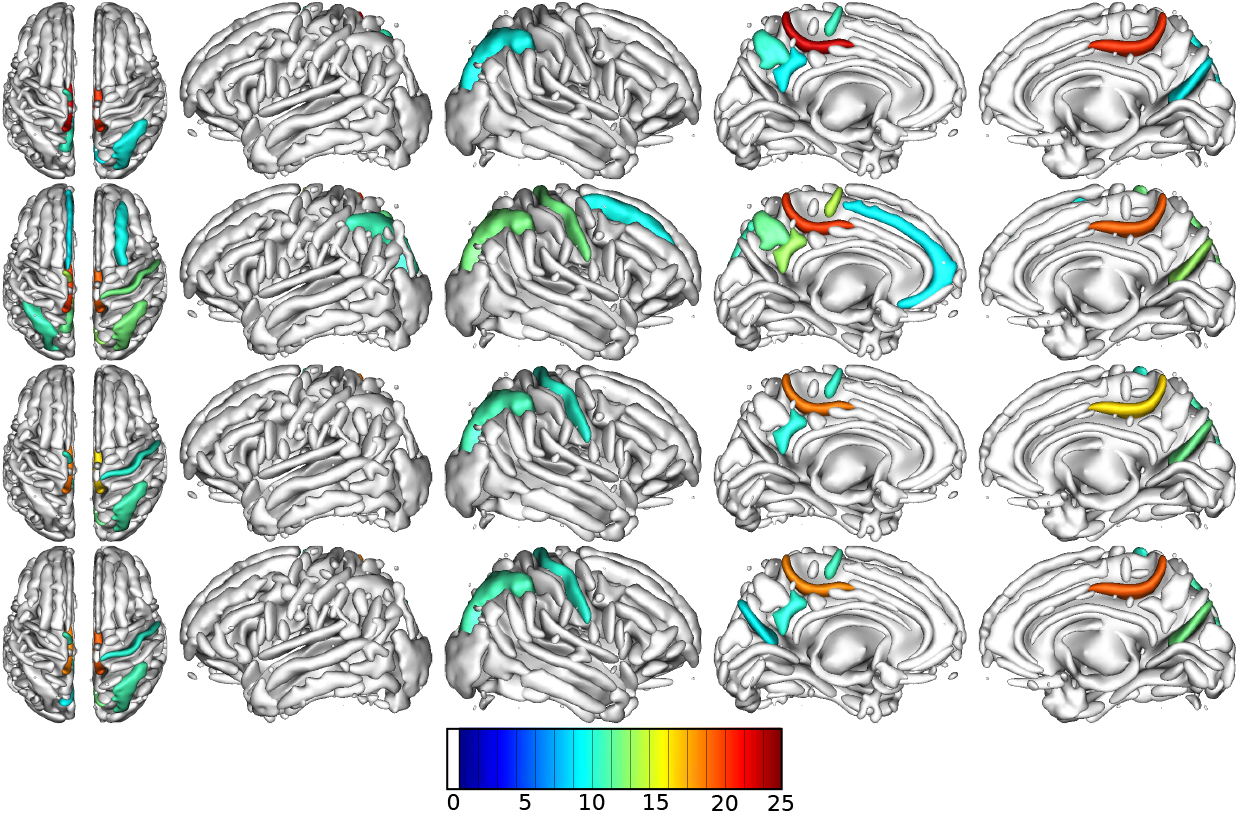
Brain sulci coloured according to –*log*10(*p*) of significant hits in the chr1:215 region. Hits - genomic region of 1 Mpb with at least one significant p-value among the three tests. In rows, from top to bottom, hits are determined using block-test, complete-test, single-test, imputed SNPs with PLINK.

**Supplementary Figure 9:**
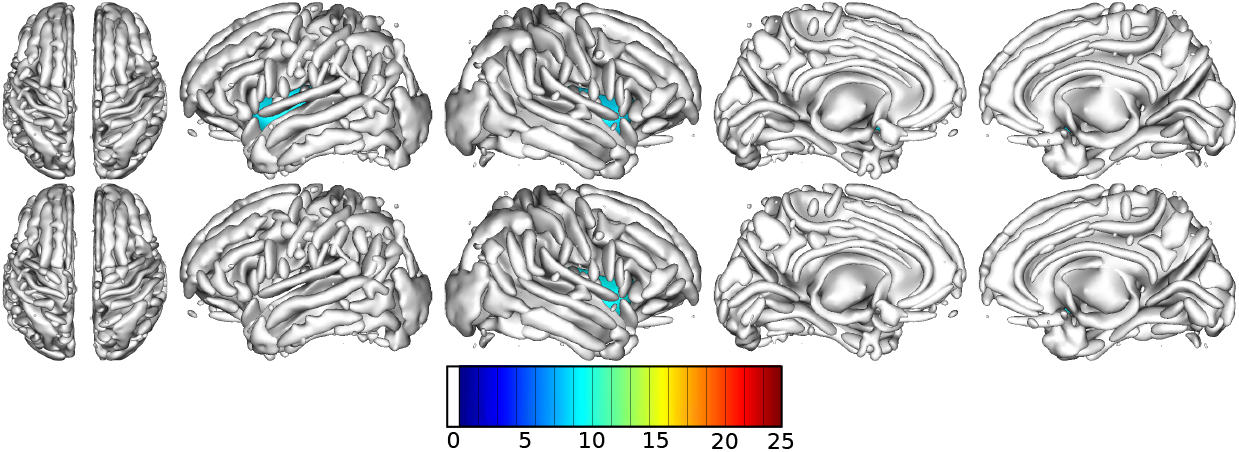
Brain sulci coloured according to –*log* 10(*p*) of significant hits in the chr7:134 region. Hits - genomic region of 1 Mpb with at least one significant p-value among the three tests. In rows, from top to bottom, hits are determined using block-test and single-test.

**Supplementary Figure 10:**
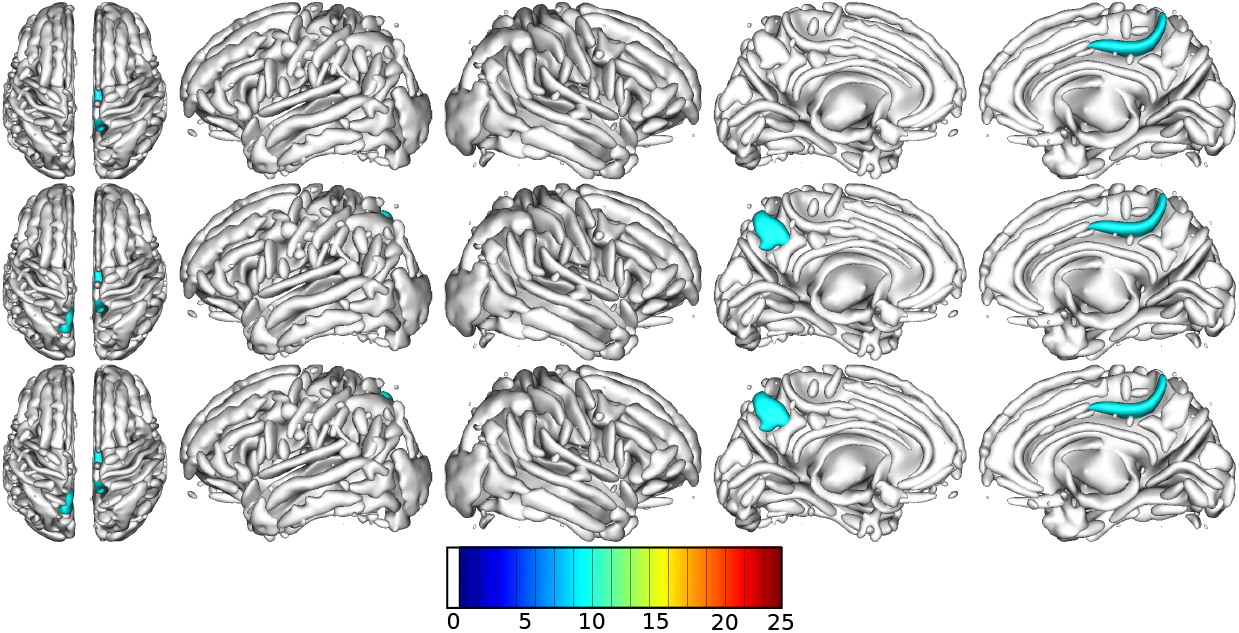
Brain sulci coloured according to –*log*10(*p*) of significant hits in the chr9:114 region. Hits - genomic region of 1 Mpb with at least one significant p-value among the three tests. In rows, from top to bottom, hits are determined using block-test, complete-test, single-test

**Supplementary Figure 11:**
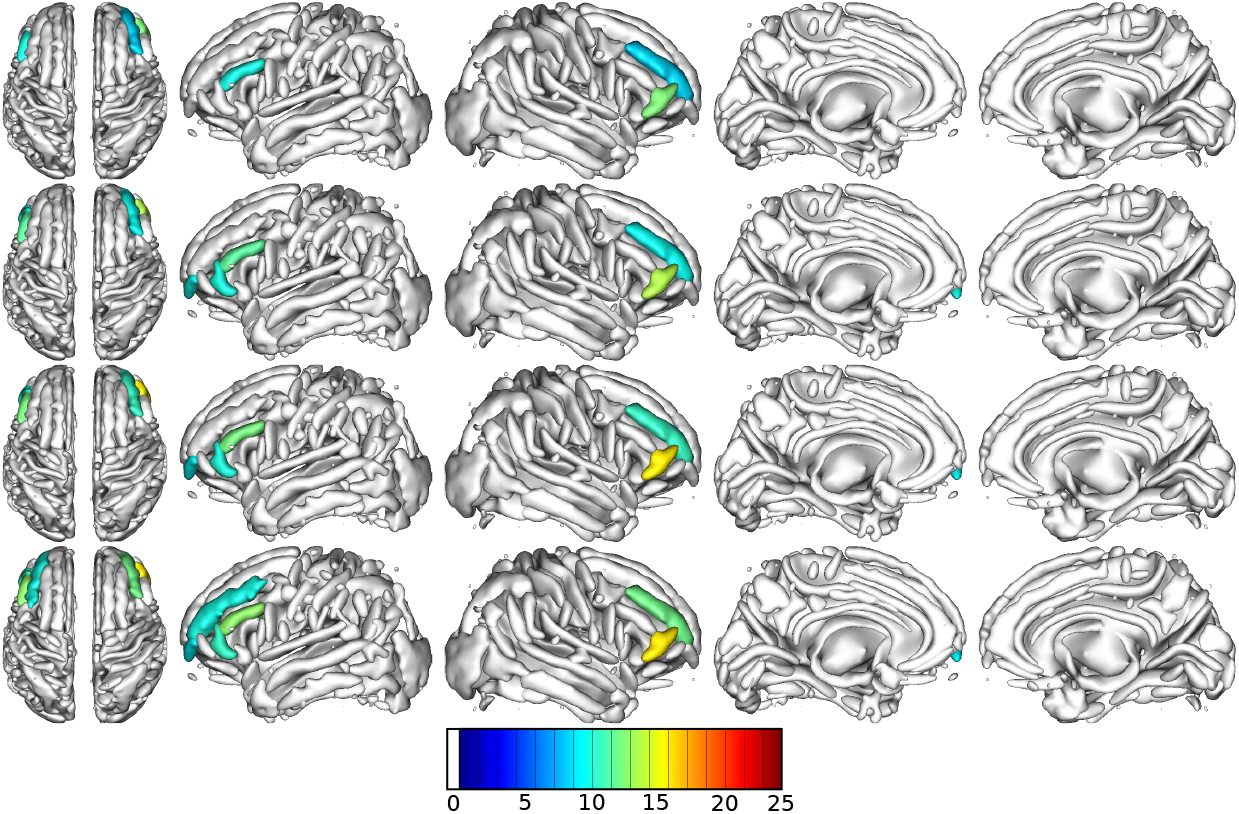
Brain sulci coloured according to –*log*10(*p*) of significant hits in the chr16:87 region. Hits - genomic region of 1 Mpb with at least one significant p-value among the three tests. In rows, from top to bottom, hits are determined using block-test, complete-test and single-test.

## Notes

### Competing Interest Statement

The authors have declared no competing interest.

